# Automatic Classification of Normal and Abnormal Cell Division Using Deep Learning Analysis of Mitosis Videos

**DOI:** 10.1101/2023.04.27.538650

**Authors:** Pablo Delgado-Rodriguez, Rodrigo Morales Sánchez, Elouan Rouméas-Noël, François Paris, Arrate Munoz-Barrutia

## Abstract

In recent years, there has been a surge in the development of methods for cell segmentation and tracking, with initiatives such as the Cell Tracking Challenge driving progress in the field. Most studies focus on regular cell population videos in which cells are segmented, cell tracks followed, and parental relationships annotated. However, DNA damage induced by genotoxic drugs or ionizing radiation provide additional abnormal cellular events of interest since they lead to aberrant behaviors such as abnormal cell divisions (i.e., resulting in a number of daughter cells different from two) and cell death.

The dynamic development of those abnormal events can be followed using time lapse microscopy to be further analyzed. With this in mind, we developed an automatic mitosis classifier that categorizes small mitosis image sequences centered around a single cell as “Normal” or “Abnormal.” These mitosis sequences were extracted from videos of cell populations exposed to varying levels of radiation that affect the cell cycle’s development. Such an approach can aid in detecting, tracking, and characterizing the behavior of the entire population.

In this study, we explored several deep-learning architectures for working with 12-frame mitosis sequences. We found that a network with a ResNet50 backbone, modified to operate independently on each video frame and then combined using a Long Short-Term Memory (LSTM) layer, produced the best results in the classification (mean F1-score: 0.93 ± 0.06). In future work, we plan to integrate the mitosis classifier in a cell segmentation and tracking pipeline to build phylogenetic trees of the entire cell population after genomic stress.

**Author Summary:** In recent years, there has been a growing interest in developing methods to analyze videos of cell populations, which show how cells move and divide over time. Typically, researchers focus on developing methods to automatically identify and track individual cells and their divisions. However, exposure to anticancer drugs or radiation can cause uncommon behaviors, such as abnormal cell divisions, which are of interest to experts studying the effects of these agents on cell behavior.

To address this issue, we developed an automated tool that can determine whether a specific cell division seen in a video is normal or abnormal. We used video microscopy to capture small sequences of cell division, and then trained a deep-learning model to classify these sequences as either normal or abnormal. We found that our model achieved a high level of accuracy in this task.

Our tool has the potential to aid experts in identifying abnormal cellular events, providing insights into the effects of genotoxic agents on cell behavior. In future work, we plan to integrate our tool into more complex methods for analyzing cell population videos, which may help us better understand the impact of toxic agents on the behavior of the entire cell population.

## Introduction

Cell segmentation and tracking have gained increased attention in recent years due to the production of new and complex data involving large cell populations and diverse behaviors. Initiatives such as the Cell Tracking Challenge (1) have brought together researchers to benchmark and advance the field. While manual methods for cell tracking (such as the one proposed in (2) for glioblastoma populations) can provide valuable insights into cell behavior, it is necessary to develop automatic methods to extrapolate this tracking to new videos. This typically involves storing the cells’ location at each timestep and its relationship with cells in previous and subsequent frames, including whether the current cell is a continuation of a previous cell or a descendant resulting from cell division and whether it is the first or last frame of the particular cell track. These parameters are commonly obtained using current cell tracking algorithms, as demonstrated in the Cell Tracking Challenge Benchmarking initiative (1,3). However, different types of cell population videos can give rise to additional variables that need to be considered, for example, in the analysis of videos that capture the cell response after genomic stress, such as those resulting from the application of radiation in therapeutical doses. Radiotherapy is widely used to compromise tumor cells and inhibit their growth (4,5), causing DNA damage that leads to unusual behaviors during mitosis, when un- or mis-repaired. In such cases, it is essential to classify each mitosis in the video as normal or abnormal to measure the disruptive effects in cell toxicity. In this work, we utilized videos of glioblastoma cell populations exposed to varying radiation levels to train and test our proposed algorithm to create a generalizable method that can be applied to different videos from future experiments.

Figure 1 illustrates the different steps of normal mitosis, abnormal mitosis resulting from cell fusion, and cell death in these videos. Normal mitosis involves two arising cells re-adhering to the plate, while cell fusion results in a single multinucleated cell. Cell populations were recorded using a contrast phase channel to show general cell shape and a far-red fluorescent channel to visualize cell nucleus after counterstaining with Sir-DNA. To automatically analyze those videos, it is necessary to identify whether a given cell division (mitosis) is normal or abnormal, which could indicate that ionizing trough DNA damage has pertubed the cell cycle’s normal functioning. For non-experts, distinguishing between normal and abnormal mitosis can sometimes be challenging due to the various ways cell division can fail.

**Figure 1:**
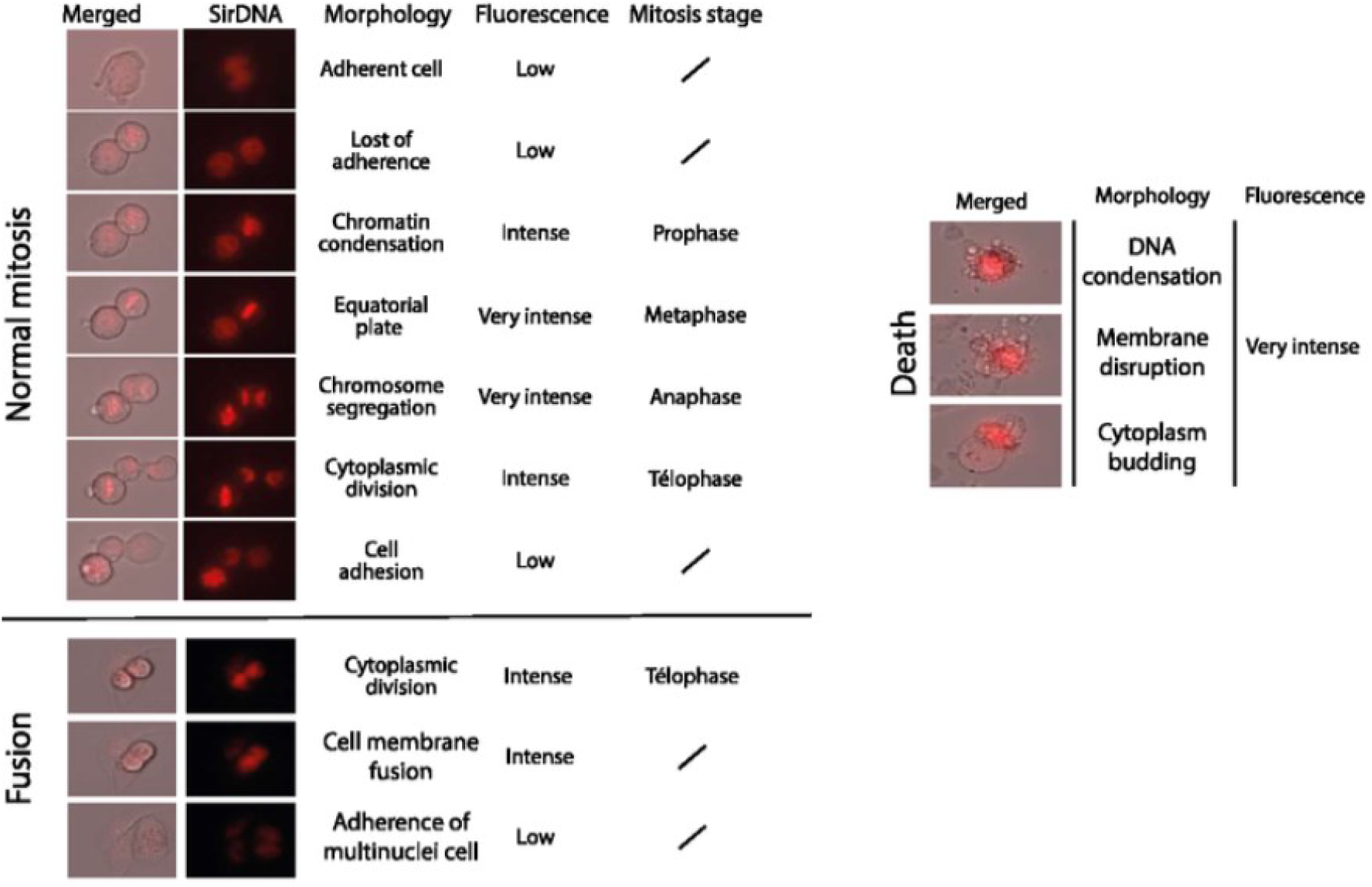
Illustration of: (Top row, left) Normal mitosis; (Bottom row, left) Fusion; (Right) Cell death. (Columns from left to right) Representative merged images (phase contrast microscopy and SirDNA); SirDNA fluorescence microscopy images; description of the cell morphology, the fluorescence level and for the normal mitoseis and the fusion, the mitosis stage. The mitosis stages show how the cell starts adhered to the plate, it detaches, the chromatin gets condensed forming a plate in the middle of the cell body, then the chromosomes are pulled to opposite borders of the cell and the cytoplasm divides. After this, both daughter cells attach again to the plate. On the contrary, in fusion, after the cytoplasm has divided it remerges again into a single cell that then attaches back to the plate. In cell death, on the other hand, the genetic material condenses, the membrane breaks and bubbles are formed.

**Figure 2:**
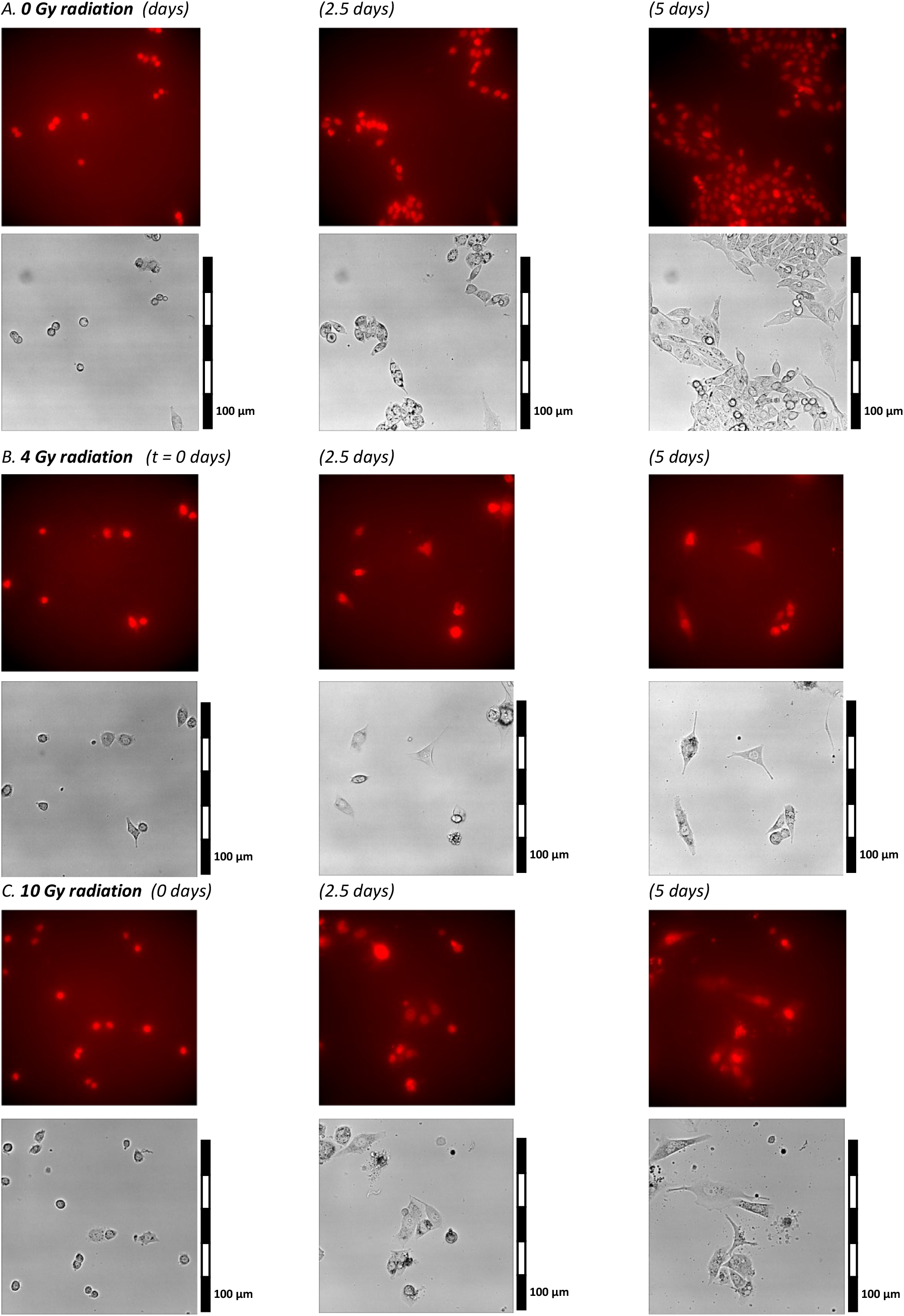
Example of cell population videos with different levels of radiation. Both channels are shown (fluorescence on top, phase contrast at the bottom). The cell populations are shown at three timesteps: 0, 2.5, and 5 days. **A**. Video with no radiation; **B**. Video with 4 Gy radiation. **C**. Video with 10 Gy radiation. Cells present more abhorrent behaviors as radiation increases, also reducing their replications (mitoses).

Most automated methods in the literature focus on detecting normal mitosis in 2D images, primarily in histology slices, to assess tumor growth (6–10). Other studies address the evolution of this process through time (11–13) to track cell division in time-lapse videos, either classifying image crops as mitosis or non-mitosis (14) or applying general classification to single-cell images (15). These studies do not consider other cell behaviors than normal mitosis during division.

Cell division occurs over a series of time steps, and the distinction between normal and abnormal mitosis is only apparent when considering temporal progression. Hence, relying on a single time frame to determine abnormality is unreliable as the entire division process must be observed. Therefore, we propose the development of an automatic classifier to identify normal or abnormal cell division in small mitosis videos centered around the dividing cell. As previously indicated, these mitosis sequences were extracted from videos of cell populations exposed to varying levels of radiation that affect the cell cycle’s development.

Deep learning algorithms have proven effective for classifying single-cell 2D images (16). We, therefore, explored several deep-learning architectures for working with our 12-frame mitosis sequences. We finally optimized a deep learning architecture with the best hyperparameters to accurately classify single-cell division videos into normal and abnormal mitosis categories. The tool could be further integrated into a cell segmentation and tracking pipeline to characterize the behavior of the entire population.

## Materials and methods

### Materials

The cells used were human glioblastoma (U251) cell lines, cultured in DMEM supplemented with 10% SVF, 1% penicillin-streptomycin, and 1% L-glycine under standard conditions at 37°C/5% CO2. Cell irradiation was performed with a CP-160 X-ray irradiator (Faxitron, Lincolnshire, Illinois, USA) at an accelerating voltage of 160 kV and a dose of 1.3Gy/min. The cells were filmed for 5 days with the Nikon Ti inverted optical microscope (Nikon, Minato-ku, Tokyo, Japan).

The initial data obtained from these cultures consisted of a series of 2D videos showing entire cell populations developing over time under different radiation levels. Each of the videos had 721 frames, 1024 × 1024 pixels per frame, and two channels, one of them showing a phase contrast image and the other one a fluorescence signal, marking the genetic material inside the cell nuclei. The images were taken at an interval of 10 min during these 5 days. 2 to 4 captures per well were made in brightfield and far-red fluorescence, which allowed the acquisition of cell morphology and nuclear labeling by SiR-Hoechst (SiR-DNA, Spirochrome).

Figure 1 shows three of the frames of these videos at 0, 2.5, and 5 days. This is done for different levels of radiation (0 Gy, 4 Gy, and 10 Gy), presenting the phase contrast and the fluorescence channels separately. It can be observed that the largest number of cell divisions, resulting in a larger cell population, occurs for 0 Gy, and that an increase in radiation causes the cells some difficulties in performing their division cycle in a normal way.

From these complete videos, several smaller image sequences were manually extracted, each centered around a mitosis event. These sequences were created by drawing a square around the mitosis event and extending it for 12 frames to ensure that each frame showed both daughter cells completely. This, in turn, corresponds to a period of two hours. The resulting sequence was stored as a separate image of 12 frames, preserving both channels, with frame sizes ranging from 50×50 to 110×110 pixels. The substack started just before cell division when the parent cell had already detached from the plate and became round and ended when both daughter cells had attached to the plate. Twelve frames were selected as the optimal length for a mitosis video as it allowed the entire division process to be observed while maintaining a manageable data size. Forty-seven substacks of each class (normal and abnormal mitosis) were stored as test data, while the remaining substacks were used for training the algorithms.

We revised all the substacks and eliminated those ambiguous for human classification as normal or abnormal to train our models to replicate human behavior. The following proportion of images was used for training and test sets: the test set would be used to evaluate the final chosen model:

- Train set
  - Abnormal mitosis: 146 substacks
  - Normal mitosis: 317 substacks
- Test set
  - Abnormal mitosis: 47 substacks
  - Normal mitosis: 47 substacks

Figure 3 shows some frames of several mitosis representative videos to illustrate differences between normal and abnormal mitoses. Both channels (phase contrast and fluorescence) are shown separately. During normal mitosis, the cell becomes round, condensing the chromosomes in the middle region of the nucleus. Then, they divide their genetic material, which migrates to each daughter cell while the cytoplasm divides. At the end of the process, two new cells are created, which can be seen in sequences A.1 and A.2 in Figure 3. On the other hand, abnormal mitosis can show several abhorrent behaviors. In B.1, the daughter cells become intermingled and end up forming a composed cell, while in sequence B.2, the division gives rise to three cells instead of two.

**Figure 3:**
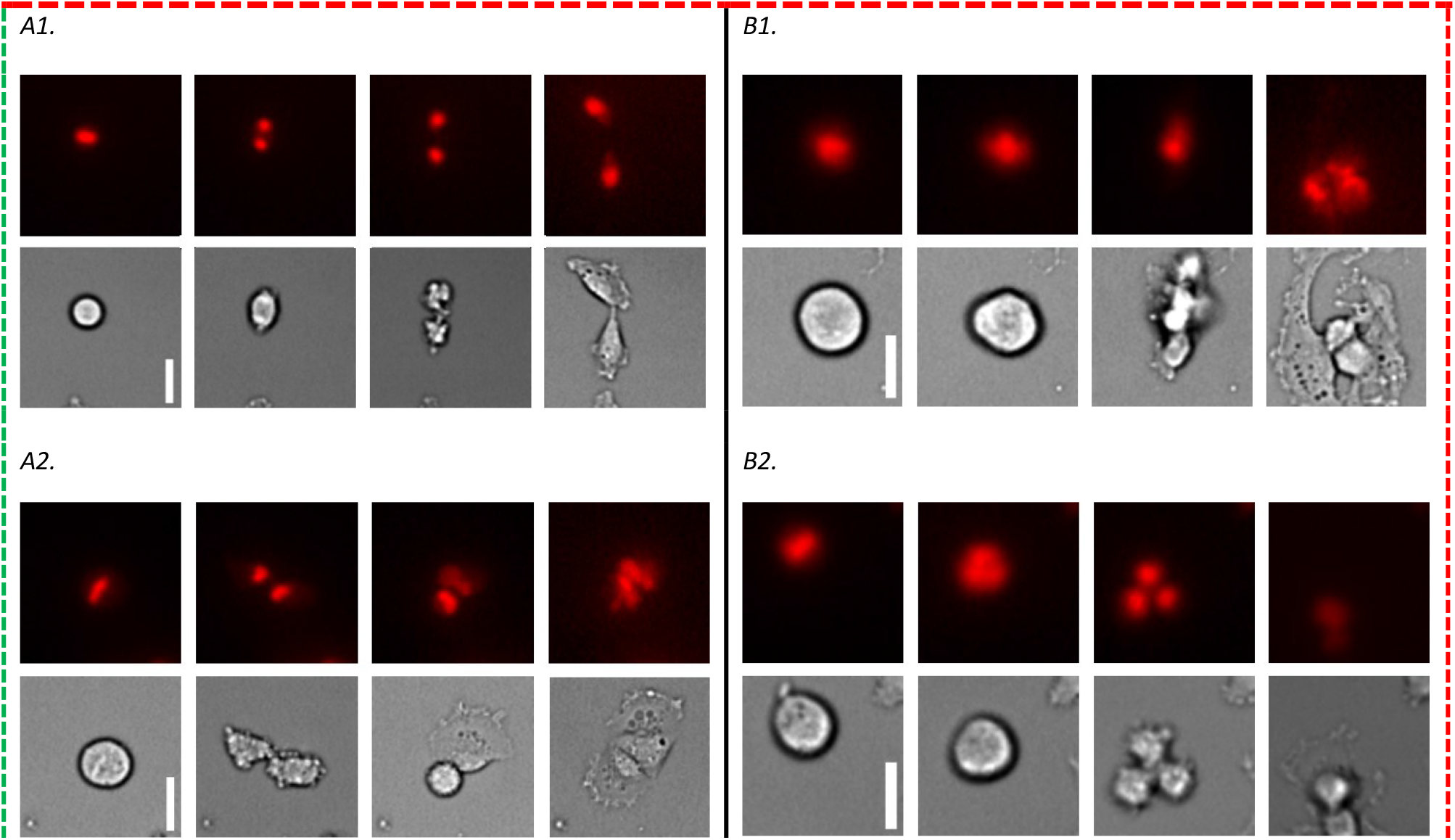
Normal vs. abnormal mitosis examples, the white scale bars represent 10 μm. Normal mitosis typically ends with two differentiated cells adhered to the plate, while abnormal mitosis may result in abnormal behaviors such as forming three cells. The image displays four frames selected from multiple mitosis substacks (original substacks consist of 12 frames, covering a period of two hours). **A1-2** Normal mitoses. **B1-2** Abnormal mitoses resulting in three cells. 4 frames are shown for each of the short videos, including the initial state, two significant middle states and the final state.

During training, we aimed to use as many images as possible, despite class imbalance. The class imbalance was addressed by weighting each class based on the proportion of images they represented. These weights were considered in the loss function. However, a balanced test set of 47 normal and abnormal mitoses was used for more reliable final score results.

### Methods

Multiple deep learning architectures were trained on the substacks to classify new samples as normal or abnormal mitosis. The temporal dimension of the samples is a unique aspect of this problem. One approach could be to treat each substack as a 3D image and use 3D convolutions, but this method would ignore the temporal relation. Instead, we used a Long Short-Term Memory (LSTM) layer in all architectures to consider the temporal relationship between consecutive frames and produce a single output summarizing the entire video’s characteristics.

The overall structure of the pipeline is shown in Figure 4. A convolutional section topped each architecture, sometimes based on known architectures like Visual Geometry Group (VGG)(17), Xception(18), or Residual Network (ResNet) (19), with the last Fully Connected (FC) layers removed. The convolutional section was applied to each frame using a “time distributed” layer from TensorFlow, with only one set of weights optimized for all frames. The output from each frame was vectorized and fed into a Long Short-Term Memory (LSTM) layer to produce a single summary of the entire video’s information. Finally, FC layers and a SoftMax layer were used to obtain the final classification output.

**Figure 4:**
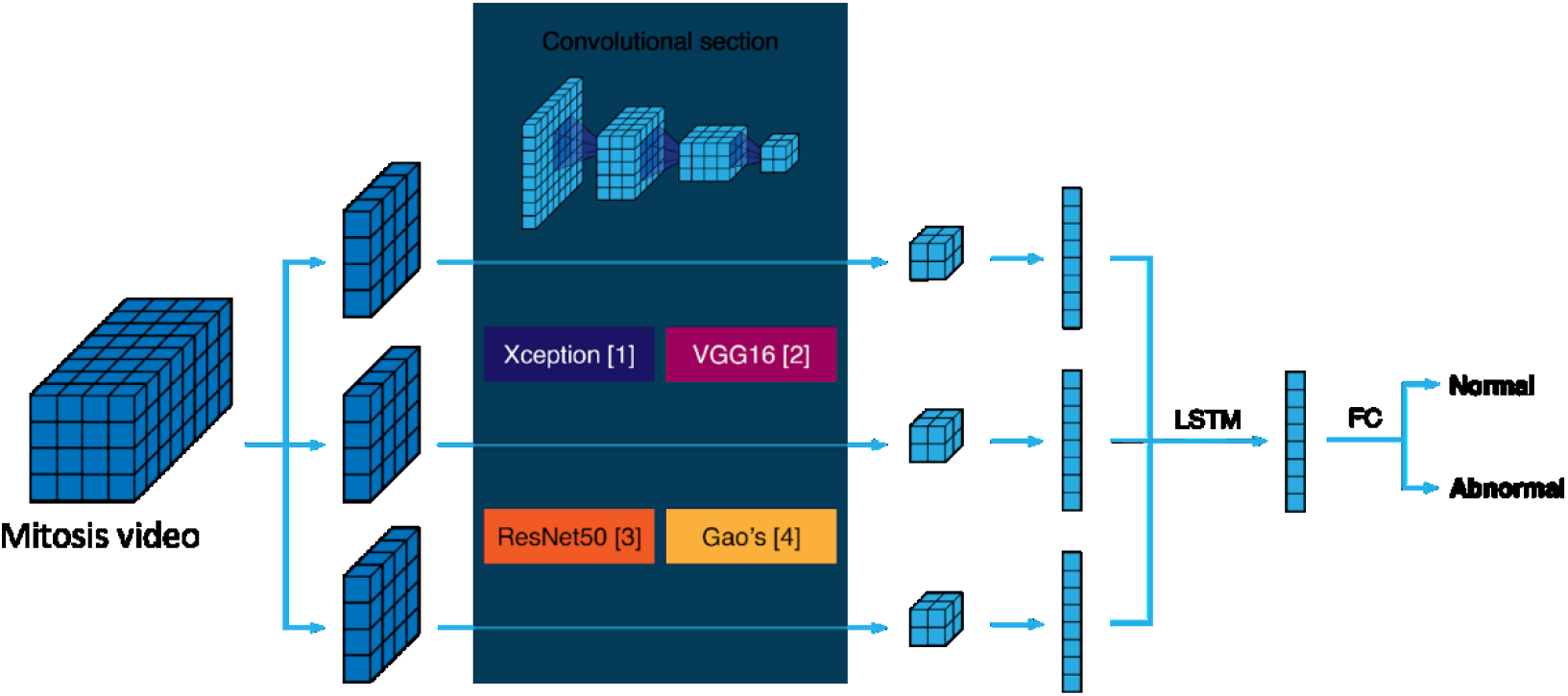
Diagram of network architectures. For each mitosis video frame, it goes through the same set of convolutional layers (Xception, VGG16, Resnet50 or Gao’s architectures) and is reduced to a feature vector. These vectors are then input to an LSTM layer to capture the temporal relationships and produce an output vector. The output vector goes through FC layers and finally yields a normal or abnormal classification.

The convolutional section of the architectures followed these structures:

- VGG16 backbone: Structured based on the use of 3×3 convolutions, smaller than its predecessors, as well as 1×1 convolutions to introduce more non-linearity.
- ResNet50 backbone: Defined for its use of residual layers that include skip connections.
- Xception backbone: Based on the Inception network (20), this architecture uses depthwise separable convolutions.
- Gao’s model, from (15): A series of layers initially used for cell classification.

To enhance the algorithms’ ability to handle diverse data and prevent overfitting, several data augmentation operations were applied to each video substack prior to the training process. These operations were performed on each frame individually, to produce a consistent outcome. The following random processes were carried out:

- Vertical flip: Randomly applied/ not applied.
- Horizontal flip: Randomly applied/ not applied.
- Rotation: Random angle within the range of 0 - 90°.
- Noise addition: Random noise added (or not) to the image, following uniform distributions for each pixel.
- Elastic deformation: A 3 × 3 pixels grid of nodes is placed overlapping 9 uniformly separated pixels of the image. These nodes are randomly deformed following uniform distributions, changing their location. At the same time, the rest of the image pixels are interpolated to follow these deformations, creating a new version of the image.

Multiple hyperparameter configurations were tested for each proposed architecture to find the optimal one and achieve the best classification. Each network was trained with each configuration 10 times, yielding a mean and standard deviation of their performance for a more reliable evaluation. The training was capped at 500 epochs with a patience of 100 epochs, meaning the training would stop if validation accuracy did not significantly improve in the last 100 epochs. The validation accuracy was obtained by evaluating the model on 43% of the randomly split training data, not used to optimize the network weights (validation data). This high validation proportion was chosen to ensure the model’s generalizability to new data. After each training, the weights with the highest validation accuracy were retained as the “best” weights.

A key parameter to analyze was the impact of using a pretrained network and fine-tuning it with our data on the classification results, compared to training from scratch with only our data. The pretrained networks were trained on ImageNet, a diverse image dataset without microscopy images. The hypothesis was that the ImageNet-trained feature extractors could aid in analyzing our images.

The network training was conducted using Google Colab pro (21), with GPU access to accelerate the computations.

## Results and Discussion

For each hyperparameter configuration, we performed ten identical pieces of training. The best validation accuracy, precision, recall, and F1-score were recorded from each of them. The mean and standard deviation for each score from the ten training runs are displayed in Table 1. This setup enabled us to obtain statistical measures of the performance of each configuration, as the stochastic optimization algorithm used in the process could choose different paths during each training run.

**Table 1:** Validation results for various architecture and hyperparameter configurations are displayed. Different colors (blue, yellow, orange, and green) indicate different architectures, with the best-performing model for each highlighted using a darker background. Each training run was repeated 10 times, and the best validation accuracy, precision, recall, and F1-score were averaged to produce a mean ± standard deviation value for each. The best-performing models were selected primarily based on the F1-score.

| Architecture | Varying hyperparameters |  |  | Results on the validation set |  |  |  |
| --- | --- | --- | --- | --- | --- | --- | --- |
|  | Pretrained | Initial learning rate | Batch size | Mean val. accuracy | Mean val. precision | Mean val. Recall | Mean val. F1-score |
| Xception | Yes | 0.001/ 0.00001 | 10 | 0.9 ± 0.03 | 0.82 ± 0.07 | 0.76 ± 0.15 | 0.77 ± 0.09 |
| Xception | No | 0.001 | 10 | 0.92 ± 0.02 | 0.89 ± 0.07 | 0.77 ± 0.07 | 0.82 ± 0.05 |
| Xception | No | 0.0001 | 10 | 0.96 ± 0.03 | 0.98 ± 0.02 | 0.86 ± 0.12 | 0.91 ± 0.07 |
| Xception | No | 0.00001 | 10 | 0.93 ± 0.04 | 0.88 ± 0.07 | 0.85 ± 0.07 | 0.86 ± 0.08 |
| Xception | No | 0.0001 | 20 | 0.94 ± 0.05 | 0.94 ± 0.06 | 0.8 ± 0.18 | 0.86 ± 0.13 |
| VGG16 | Yes | 0.001/ 0.00001 | 10 | 0.94 ± 0.02 | 0.92 ± 0.06 | 0.82 ± 0.13 | 0.86 ± 0.07 |
| VGG16 | No | 0.001 | 10 | 0.75 ± 0 | 0 ± 0 | 0 ± 0 | 0 ± 0 |
| VGG16 | Yes | 0.0001/ 0.00001 | 10 | 0.75 ± 0 | 0 ± 0 | 0 ± 0 | 0 ± 0 |
| VGG16 | Yes | 0.00001/ 0.000001 | 10 | 0.94 ± 0.02 | 0.91 ± 0.07 | 0.84 ± 0.11 | 0.86 ± 0.05 |
| VGG16 | Yes | 0.00001/ 0.000001 | 20 | 0.93 ± 0.04 | 0.9 ± 0.1 | 0.83 ± 0.1 | 0.86 ± 0.09 |
| ResNet50 | Yes | 0.001/ 0.00001 | 10 | 0.95 ± 0.04 | 0.89 ± 0.1 | 0.91 ± 0.1 | 0.9 ± 0.09 |
| ResNet50 | No | 0.001 | 10 | 0.82 ± 0.04 | 0.66 ± 0.1 | 0.53 ± 0.14 | 0.58 ± 0.11 |
| ResNet50 | Yes | 0.0001/ 0.00001 | 10 | 0.94 ± 0.05 | 0.93 ± 0.08 | 0.81 ± 0.17 | 0.86 ± 0.13 |
| ResNet50 | Yes | 0.00001/ 0.000001 | 10 | 0.95 ± 0.04 | 0.91 ± 0.1 | 0.89 ± 0.09 | 0.9 ± 0.09 |
| ResNet50 | Yes | 0.001/ 0.00001 | 20 | 0.85 ± 0.05 | 0.8 ± 0.11 | 0.51 ± 0.15 | 0.62 ± 0.14 |
| Gao | No | 0.001 | 10 | 0.86 ± 0.05 | 0.74 ± 0.1 | 0.69 ± 0.2 | 0.7 ± 0.15 |
| Gao | No | 0.0001 | 10 | 0.83 ± 0.07 | 0.65 ± 0.13 | 0.75 ± 0.11 | 0.69 ± 0.1 |
| Gao | No | 0.00001 | 10 | 0.72 ± 0.03 | 0.33 ± 0.18 | 0.31 ± 0.2 | 0.31 ± 0.18 |
| Gao | No | 0.001 | 20 | 0.84 ± 0.04 | 0.66 ± 0.24 | 0.58 ± 0.21 | 0.61 ± 0.21 |

The best-performing model on validation data from each architecture was selected. The ten equally trained models corresponding to that parameter setup were evaluated on the test set to determine their final performance on unseen data. Table 2 displays the test results for these best-performing models. This final evaluation step is necessary to obtain a reliable measure of the network’s overall performance and allows us to rank these four alternatives. The F1-score results, used as an overall score, exceed 0.79, indicating all the algorithms performed well, with some exhibiting particularly reliable behavior.

**Table 2:** Test scores for the best-performing model of each architecture. The model with the highest mean F1-score and lowest standard deviation is highlighted with a grey border.

| Architecture | Varying hyperparameters |  |  | Results on the test set |  |  |  |
| --- | --- | --- | --- | --- | --- | --- | --- |
|  | Pretrained | Initial lr | Batch size | Mean accuracy | Mean precision | Mean Recall | Mean F1-score |
| Xception | No | 0.0001 | 10 | $0.94 \pm 0.07$ | $1 \pm 0.01$ | $0.88 \pm 0.14$ | $0.93 \pm 0.1$ |
| VGG16 | Yes | 0.00001/<br>0.000001 | 10 | $0.89 \pm 0.05$ | $0.95 \pm 0.03$ | $0.83 \pm 0.11$ | $0.88 \pm 0.06$ |
| resnet50 | Yes | 0.001/<br>0.00001 | 10 | $0.94 \pm 0.05$ | $0.97 \pm 0.03$ | $0.9 \pm 0.09$ | $0.93 \pm 0.06$ |
| Gao | No | 0.001 | 10 | $0.82 \pm 0.08$ | $0.93 \pm 0.03$ | $0.71 \pm 0.16$ | $0.79 \pm 0.12$ |

In summary, the approach presented in this article aims to automate the analysis of videos of cell populations subjected to toxicity. Tracking and segmenting these cells can provide valuable information on their behavior under different levels of toxicity. However, as mentioned earlier, cells can exhibit unusual behaviors in these cases, particularly unstable cell division, which can be challenging for the human eye to evaluate quickly. A deep neural network algorithm can classify mitoses without defining and measuring specific cell characteristics, as no clear parameters solely define normal or abnormal mitosis. Our algorithm, therefore, addresses the need for an automatic tool to tackle this specific classification task and can be integrated with tracking and segmentation tools in the future.

One of the project objectives was to assess the impact of using a pretrained network on mitosis classification results, even if the pretraining dataset (ImageNet) contained very diverse images unrelated to microscopy. Only networks with widely used backbones (ResNet, VGG, or Xception) could be pretrained. As expected, the pretrained versions of VGG16 and ResNet50 performed better. The hypothesis was that, despite being trained on single (non-video) images of various types, the network would have learned useful feature extractors that would improve results compared to starting from scratch with our data. Nevertheless, the VGG16 architecture showed better average results when its weights were randomly initialized instead of pretrained. This could be because this architecture is more capable of working directly with our images and can find optimal values for its weights more easily, or because the random initializations were fortunate and set up the model for better results.

Regarding the initial learning rate of the optimizer, both Xception and VGG16 architectures performed best on the validation set with a lower value. VGG16, in particular, offered no true positives for both the configuration trained from scratch with 0.001 as initial learning rate and the pretrained configuration for an initial rate of 0.0001. This extreme behavior was mitigated by decreasing the initial learning rate to 0.00001, with a learning rate to train the last unfrozen layers at 0.000001 for a pretrained configuration, which led VGG16 to achieve optimal results. Low learning rate values improve the reliability of finding the cost function’s minimum but can also result in local minima instead of global ones. ResNet50 offered higher performance for the pretrained configuration without using very low initial learning rate values. On the other hand, the model based on Gao’s architecture performed better with a higher learning rate of 0.001, as it was not pretrained and needed larger steps to find a suitable minimum. The third hyperparameter to be tuned, batch size, did not have a significant impact. Increasing the batch size did not improve the results for any of the architectures.

Ultimately, the best versions of ResNet50 (pretrained) and Xception (trained from scratch) outperformed all other models on the test set, achieving a mean F1-score of 0.93 on test data. ResNet50 had a lower standard deviation, making it the best model overall after considering high individual scores for accuracy, precision, and recall. Surprisingly, a trained-from-scratch Xception model performed comparably to a ResNet50 model with prior knowledge. The chosen top-performing algorithm, ResNet50, provides accurate results for the classification problem and can support human human experts in the classification task.

Figure 5 shows examples of mitoses misclassified by the ResNet50 model. These errors could be attributed to several factors, as the network can be easily misled by various events, including overlapping cells, crowded surroundings, poor contrast in some videos, and instances where the daughter cells do not have the time to re-attach to the surface during the video, as seen on example B2. It can be concluded that the algorithmic pipeline is particularly adept at identifying mitosis as normal or abnormal when cells are not too crowded together and in the cases (most of them) where a selection of 12 frames (50 min) covers the whole mitotic process.

**Figure 5:**
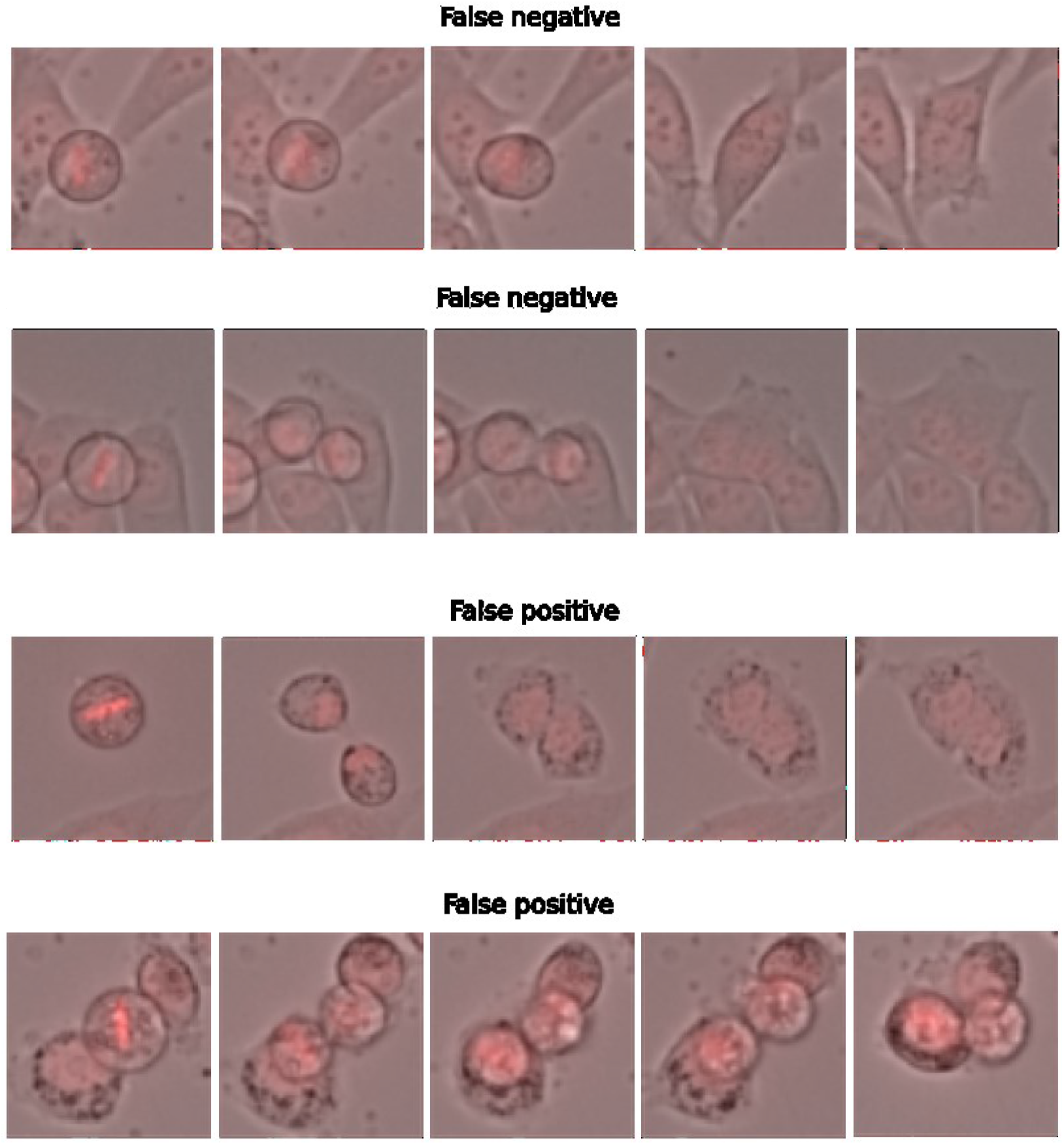
Examples of false negatives (abnormal mitosis classified as normal) and false positives (normal mitosis classified as abnormal) from the ResNet50 model applied to test data. Each example displays 4 of the 12 frames in each mitosis sample. In **A1**, a cell clearly divides completely and then reunites, making the mitosis abnormal, but the algorithm might mistake the initial division as normal. In **A2**, the cell tries to divide but fails, leading the algorithm to mistake a different cell in the image as a second daughter and classify the mitosis as normal. In **B1**, the cell divides correctly, but a different cell appearing in the image might confuse the network into misclassifying the mitosis as abnormal. In **B2**, both daughter cells end up separate, but the algorithm might mistake them as one cell, leading to classification failure.

## Conclusion

In this study, our goal was to simplify the task of analysing cell mitosis by developing an automatic method for classifying mitotic events as normal or abnormal. After evaluating different approaches and configurations, our results demonstrated that a neural network with a ResNet50 backbone and an LSTM layer accurately classified small mitosis video sequences, achieving a mean F1-score of 0.93 ± 0.06. Interestingly, we found that pretraining on ImageNet may not be essential for this type of image analysis, as a similar score was obtained using an Xception backbone without pretraining. This finding suggests that the Xception architecture might be particularly effective for analyzing microscopy data.

Given the growing importance of cell segmentation and tracking algorithms in characterizing the cellular behavior within specific environments, the classifier presented here can streamline a tedious task and be integrated into a larger workflow, such as a mitosis detection system, to identify the locations of normal and abnormal mitoses throughout a cell population over time. In the future, this approach could enable a comprehensive understanding of the cell population’s behavior under radiation or other genomic stresses. It could be used to observe the effects of these treatments on commonly-used 2D cultures from any adherent cancer cell types, such as glioblastoma shown in this present work. Additionally, the algorithm could potentially be expanded to discriminate the different types of abnormal mitosis, including fusion, non-dichotomic division (in three or more cells) or loss of genomic materials in micronuclei. All those events characteristic of genomic instability lead to shuttle genome of tumor cells.

Furthermore, the proposed method can be applied to microscopy images obtained from different studies investigating the effects of various toxic chemical compounds on normal cell cycle development. The key aspect to consider is that this algorithm can be retrained with the appropriate set of images, allowing researchers to automate mitosis classification tasks and characterize the development of cell populations exposed to toxic chemical compounds or other forms of toxicity.

## Data availability

A version of the code has been made available as a Google Colab notebook in the style of ZeroCostDL4Mic (22), aiming to facilitate its reusability by not-programming users. This notebook allows training the different architectures on a set of mitosis videos (12 frames each) specified by the user and is available at (23). This code uses a version of the github video generator repository (24). The performance of the trained network can also be tested on a different data set, and there is also an option to apply the algorithm to new data to obtain predictions. All the instructions to perform these operations are clearly explained in the notebook and the train and test mitosis videos, as well as the hyperparameters stored from the trainings, can be found at (25). The user can run all the processes on the Google servers, so no specific hardware is required. The set of short mitosis videos used during the experiments is also provided, and it is possible to train and test the networks on them.

## Acknowledgments

Special thanks to Irene Rubia-Rodríguez for designing the network schematics figure and the tables.

Thanks also to Renato Antoniassi, Margaux Crédeville, Juan-David Garcia, Abdelhadi Harraq and Benjamin Heinrich, students from the École Centrale de Nantes, for their help with the algorithms.

This work was partially funded by Ministerio de Ciencia, Innovación y Universidades, Agencia Estatal de Investigación, under Grants FPU19/02854 and PID2019-109820RB-I00, MCIN/AEI/10.13039/501100011033/, co-financed by European Regional Development Fund (ERDF), “A way of making Europe”. A. M. B. acknowledges support from Universidad Carlos III de Madrid (“Ayudas para la recualificación del profesorado funcionario o contratado”).

